# The wheat multi-pathogen resistance gene, *Lr67res*, confers a novel gain-of-function phenotype involving anion fluxes

**DOI:** 10.1101/2022.03.22.485337

**Authors:** Ricky J Milne, Katherine E Dibley, Jayakumar Bose, Adnan Riaz, Jianping Zhang, Wendelin Schnippenkoetter, Anthony R Ashton, Peter R Ryan, Stephen D Tyerman, Evans S Lagudah

## Abstract

Partial resistance to multiple biotrophic fungal pathogens in wheat (*Triticum aestivum* L.) is conferred by the *Lr67* gene, which encodes a <u>S</u>ugar <u>T</u>ransport <u>P</u>rotein 13 (STP13) family hexose-proton symporter variant. Two mutations (G144R, V387L) differentiate the resistant and susceptible protein variants (Lr67res and Lr67sus). The molecular function of the Lr67res protein is not well understood. We functionally characterized the wheat Lr67res protein variant using two heterologous expression systems – *Xenopus laevis* oocytes and *Saccharomyces cerevisiae* yeast. Wheat and barley (*Hordeum vulgare*) were used to verify disease resistance capability of Lr67/STP13 variants. The *Lr67res* allele, but not *Lr67sus*, induced large sugar-independent, anion-dominated currents in oocytes and an increased sensitivity to ions in yeast, implicating a novel gain-of-function. We demonstrate that the single mutant variant, Lr67sus^G144R^, confers disease resistance in wheat and that transgenic barley (*Hordeum vulgare* L.) plants expressing the orthologous *HvSTP13* gene carrying the equivalent mutations present in *Lr67res* exhibited increased resistance to *Puccinia hordei*. NaCl treatment was found to induce leaf tip necrosis in *Lr67res* wheat. An Lr67res-like gain-of-function can be introduced into orthologous plant hexose transporters via single amino acid mutation, highlighting the possibility of generating disease resistance in other crop species, especially with gene editing.

**One sentence summary:** The wheat Lr67res protein responsible for multipathogen resistance has a gain-of-function over the Lr67sus protein, that is associated with anion fluxes and likely contributes to disease resistance.

## INTRODUCTION

Wheat is the major crop grown throughout temperate regions of the world. Many wheat cultivars are susceptible to the disease-causing pathogens prevalent in these regions. Pathogen infection is the leading biotic cause of grain yield losses, with cumulative global losses due to the three wheat rust diseases − stem rust, leaf rust, and yellow/stripe rust – reaching around US$2.9 billion in value annually (Huerta-Espino et al., 2020). The fungal pathogens responsible for these diseases (*Puccinia graminis* f. sp. *tritici, P. triticina* and *P. striiformis* f. sp. *tritici*, respectively) siphon nutrients from infected plants which compromises growth and yield. A chemical-free strategy for controlling fungal diseases utilizes the plant’s own disease resistance genes. Strong resistance can be conferred by seedling *R* genes, which recognize pathogen effectors and trigger localized cell death, known as the hypersensitive response, to limit the spread of pathogens to other living tissues. The gene-for-gene interaction between R proteins and pathogen effectors is often pathogen race-specific and may be overcome when a pathogen effector evades R protein recognition. This type of resistance breakdown was exemplified by the emergence of Ug99, a stem rust race group highly virulent against the resistance present in many commercial wheats (Pretorius et al., 2000). There is an ongoing need to discover and introgress new R genes into elite cultivars of wheat to combat the constant evolution of fungal effector proteins. Hence sources of durable resistance are required to protect crops against fungal epidemics.

Adult plant resistance (*APR*) genes are less common than seedling *R* genes and some have the additional desirable characteristic of conferring durable resistance to multiple pathogens. Only a handful of such genes (including *Lr67, Lr34* and *Lr46*) have been described in bread wheat, compared with ∼200 race-specific seedling *R* genes. *Lr67* encodes a sugar transporter orthologous to the *Arabidopsis thaliana* Sugar Transport Protein 13 (STP13), *Lr34* encodes an ATP-binding cassette (ABC) transporter and *Lr46* remains to be fully characterized (Singh et al., 1998; Krattinger et al., 2009; Kolmer et al., 2015; Moore et al., 2015). Unlike race-specific *R* genes that rely on a gene-for-gene interaction with pathogen avirulence genes, *APR* genes confer partial resistance to multiple races of wheat rust and powdery mildew (*Blumeria graminis* f. sp. *tritici*) and are less susceptible to being overcome by the variable nature of obligate pathogen genetics (Ellis et al., 2014). Interestingly, transgenic expression of *Lr34* or *Lr67* in other cereals confers resistance to fungal pathogens adapted to those particular species (Risk et al., 2013; Krattinger et al., 2016; Schnippenkoetter et al., 2017; Sucher et al., 2017; Milne et al., 2019). The mechanism by which *APR* genes confer resistance is unclear but appears to be fundamentally different from *R* genes. In bread wheat, *APR* genes function mainly in adult plants where they slow pathogen growth without a hypersensitive response. Since *APR* genes only confer partial resistance they can be combined with NLR genes to provide a stronger and more durable resistance (Ellis et al., 2014).

This study investigated *Lr67* function. Two *Lr67* alleles occur in wheat: the common wild-type *Lr67sus* allele and the rare disease-resistance allele *Lr67res*, distinguished by two amino acid mutations – G144R and V387L. The G144 residue is highly conserved across sugar porter (SP) family members and the G144R change alone causes the protein to lose its original hexose transport function (Moore et al., 2015). Structural determination of the closely-related sugar transport protein AtSTP10 from Arabidopsis (Paulsen et al., 2019) gives additional insight into the importance of the G144 residue in proton coupling (Bavnhøj et al., 2021). Lr67res-mediated disease resistance was proposed to occur via a dominant-negative interference whereby the non-functional Lr67res protein forms inactive transporter complexes with the functional homeologs on the A and B genomes of hexaploid wheat. However, this study provides the first evidence that the APR Lr67res confers a novel gain-of-function characterized by enhanced ion fluxes that likely underpins resistance. We used *Xenopus laevis* oocytes and yeast (*Saccharomyces cerevisiae*) as heterologous expression systems as well as wheat and barley to examine function of Lr67/STP13 variants *in planta*.

## RESULTS

### Lr67sus mediates H^+^-coupled hexose transport in oocytes whereas Lr67res induces novel anion transporter-like properties

To examine functional characteristics of their encoding proteins, *Lr67sus* and *Lr67res* cRNA was injected into *X. laevis* oocytes and analyzed using electrophysiology and isotopic techniques. *Lr67sus* expression in oocytes induced transport properties typical of other proton-coupled sugar transporters (Boorer et al., 1992; Carpaneto et al., 2005; Sun et al., 2010). More specifically, *Lr67sus* oocytes displayed glucose-dependent inward currents that shifted the reversal potential from +8 mV to +33 mV and mediated greater ^14^[C]-glucose uptake than water-injected controls (Figure 1A-C). The voltage-dependent affinity of Lr67sus for glucose and its substrate specificity (Figure 1D-F) were similar to other STPs (Hayes et al., 2007; Büttner, 2010; McCurdy et al., 2010). Further, the application of a pH 4.5 glucose bathing solution to unclamped *Lr67sus* oocytes resulted in greater cytosolic acidification (Figure 2A) and a larger membrane depolarization than measured in control oocytes (Figure 2B). By contrast, *Lr67res* expression in oocytes did not increase ^14^[C]-glucose uptake (Figure 1B) nor did it cause glucose-dependent currents or shifts in reversal potential (Figure 2C). Instead, *Lr67res* expression induced large glucose-independent currents that reversed near −30 mV (Figure 2D). The magnitude of these currents was ∼10-fold greater than the glucose-dependent currents measured in *Lr67sus* oocytes (Figure 1A).

**Figure 1.**
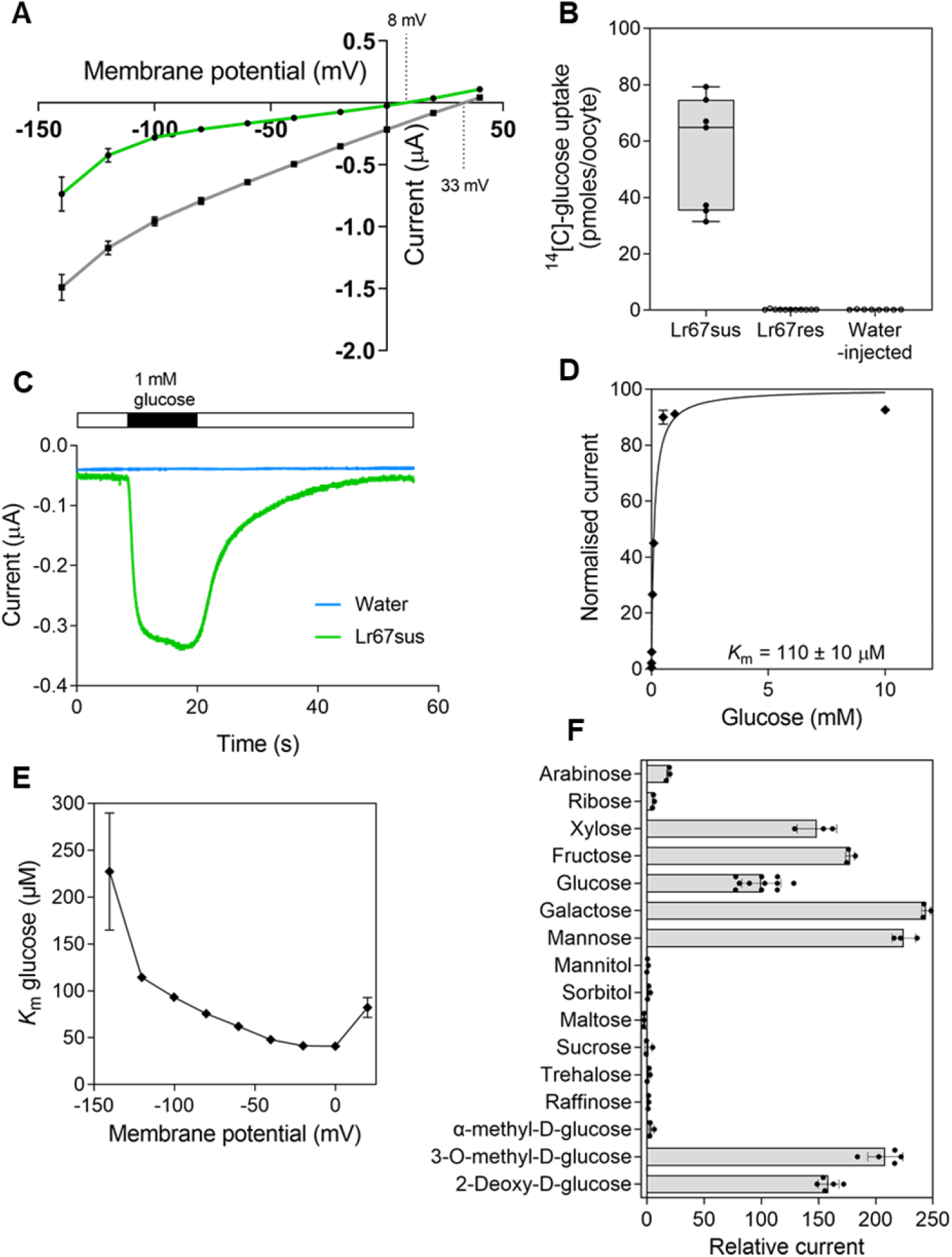
Functional characterization of Lr67sus in *X. laevis* oocytes. **A**, Current-voltage relationship of *Lr67sus*-injected oocytes under voltage-clamped conditions, perfused with 115 mM NaCl Ringer solution ± 1 mM glucose; dotted lines indicate reversal potential values. **B**, One-hour glucose uptake by oocytes injected with *Lr67sus, Lr67res* or water, incubated in 115 mM NaCl Ringer at pH 5 containing 11.4 μM ^14^[C]-glucose. **C**, Current trace of single water injected (blue) or *Lr67sus-*injected (green) oocytes clamped at –40 mV and perfused with 115 mM NaCl Ringer ± 1 mM glucose. **D**, Kinetic analysis of glucose transport by *Lr67sus-*injected oocytes under voltage-clamped conditions at a membrane potential of −120 mV. A Michaelis-Menten curve was fitted which was then normalized to *V* _max_ and plotted against the substrate concentration. **E**, Voltage dependence of Lr67sus glucose *K*_m_ at membrane potential values from –140 mV to 20 mV. **F**, Substrate selectivity of *Lr67sus* injected oocytes, currents induced at –120 mV are shown and are normalized to the current obtained for 1 mM glucose; other substrate concentrations = 5 mM. Data points with vertical/horizontal bars represent the mean ± SE of 3-11 oocytes (**A, D-F**). Boxes correspond to values within the 25th and 75th percentiles and the horizontal line represents the median; whiskers show max and min values, n=7-12 oocytes (**B**).

**Figure 2.**
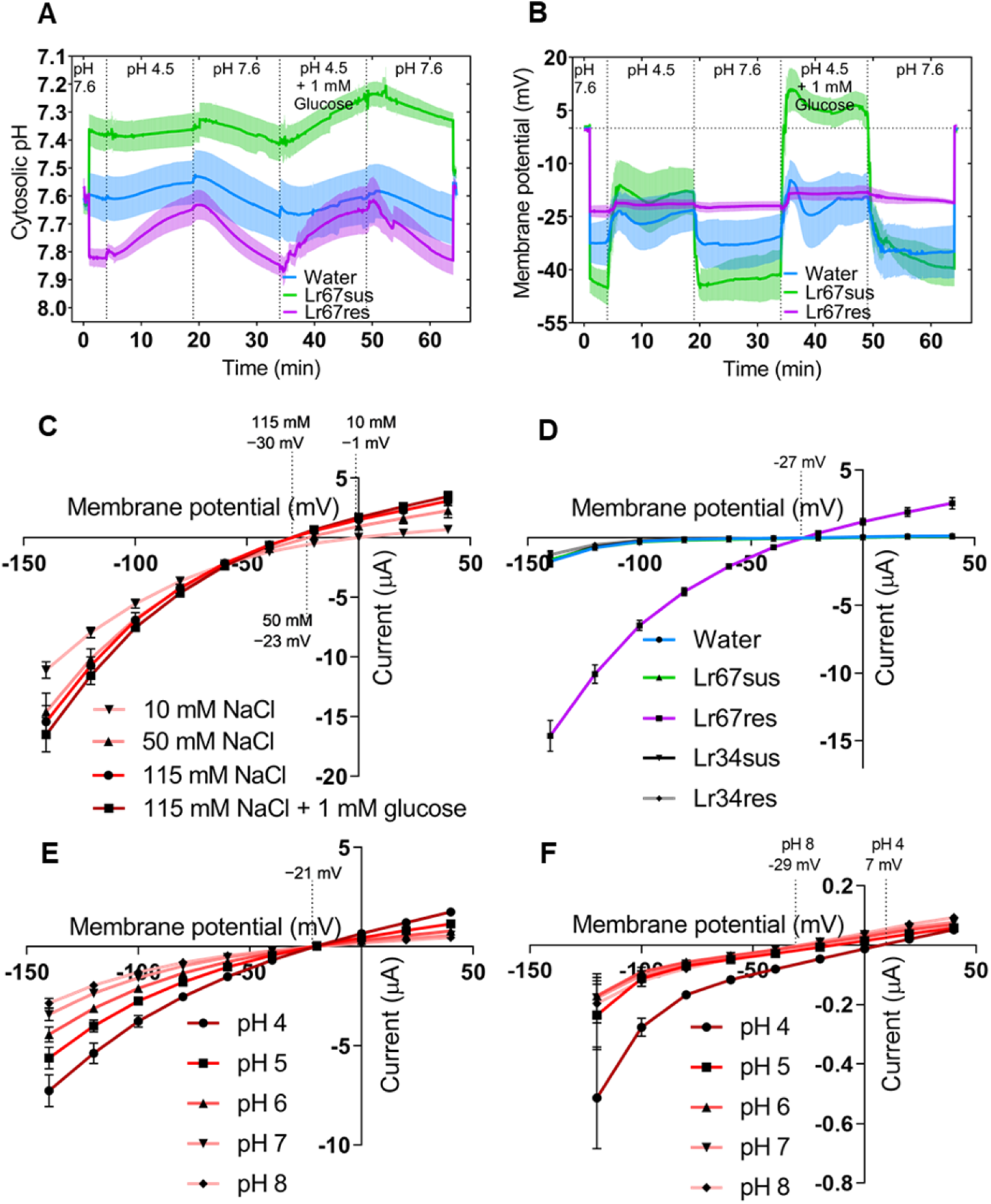
Effect of external pH and substrate concentration on clamped and unclamped oocytes. **A**, Cytosolic pH and **B**, membrane potential of unclamped *Lr67sus, Lr67res* or water-injected oocytes exposed to 115 mM NaCl Ringer solution with different pH values and glucose concentrations. Current-voltage relationship of **C**, *Lr67res*-injected oocytes perfused with NaCl Ringer solution with different concentrations of NaCl and glucose; **D**, oocytes injected with water, *Lr67sus, Lr67res, Lr34sus* or *Lr34res*, perfused with 115 mM NaCl Ringer solution; **E**, *Lr67res*-injected or **F**, water-injected oocytes perfused with 115 mM NaCl Ringer solution at pH values from pH 4 to pH 8. Dotted lines indicate reversal potential values in **C**-**F**. Dark traces with light shading represent the mean ± SE of 5–6 oocytes (**A**,**B**). Data points with vertical bars represent the mean ± SE of 4–11 oocytes (**C**-**F**).

Cytosolic pH in *Lr67res* oocytes was initially more alkaline than the water-injected controls while *Lr67sus* oocytes were more acidic, yet the changes in the proton concentration in the cytosol caused by changing the bathing solution from pH 7.6 to pH 4.5 were similar for all oocytes (Figure 2A; see Supplemental Figure S1 accounting for the logarithmic pH scale). The membrane potentials of *Lr67sus* and control oocytes showed large shifts as the pH and glucose concentrations in the bathing solution were varied, whereas the membrane potential of *Lr67res* oocytes changed little and remained near −20 mV for all treatments (Figure 2B). This response was confirmed by additional TEVC experiments showing that acidic solutions induced large increases in current magnitude in *Lr67res* oocytes without shifting the reversal potential (Figure 2E). These same treatments caused much smaller changes of current in control oocytes and shifted the reversal potential in the positive direction by 36 mV (from −29 mV at pH 8 to +7 mV at pH 4; Figure 2F). The absence of any depolarization in *Lr67res* oocytes as pH changed from pH 8 to pH 4 indicates that protons were not a major contributor to the current.

To investigate the large currents in *Lr67res* oocytes in more detail we first reduced the NaCl concentration in the bathing solution. As NaCl concentration was changed from 115 mM to 10 mM NaCl the current magnitudes decreased, and the reversal potentials shifted in a positive direction from −30 mV to −1 mV (Figure 2C). This contrasted with positive shifts in reversal potential as external NaCl was increased in oocytes injected with either the wheat or Arabidopsis *High-affinity K*^*+*^ *transporter 1* (*TaHKT1;5-A* and *AtHKT1* respectively) both of which are known to transport Na^+^ (Uozumi et al., 2000; Munns et al., 2012). The Na^+^ concentration in *Lr67res*-injected oocytes was ∼60% greater than the *Lr67sus* or water-injected oocytes (Figure 3A). This was investigated further by changing the ions in the bathing solution. Substituting Na^+^ with K^+^ or choline^+^, whilst maintaining a constant Cl^−^ concentration, caused only minor changes to the current magnitudes and reversal potentials. By contrast, substituting Cl^−^ for NO_3_^−^ or gluconate^−^ caused much larger changes to both the current magnitudes and reversal potentials (Figure 3B). Substituting 115 mM NaCl with 115 mM NaNO_3_ increased the inward and outward currents and shifted the reversal potential from −25 mV to −42 mV, which is toward the predicted reversal potential for NO_3_^−^, given an estimated internal concentration of <1 mM in oocytes (Liu et al., 1999; Liu and Tsay, 2003). Substitution with the less-permeable gluconate^−^ anion reduced the current magnitude and shifted the reversal potential in the positive direction by 51 mV from −26 mV to +26 mV (Figure 3B). NO_3_^−^ and gluconate^−^ were included in these experiments because, in heterologous expression systems, many plant anion channels are permeable to anions that may not necessarily be their primary substrate *in planta* (Kollist et al., 2011; Wang et al., 2012).

**Figure 3.**
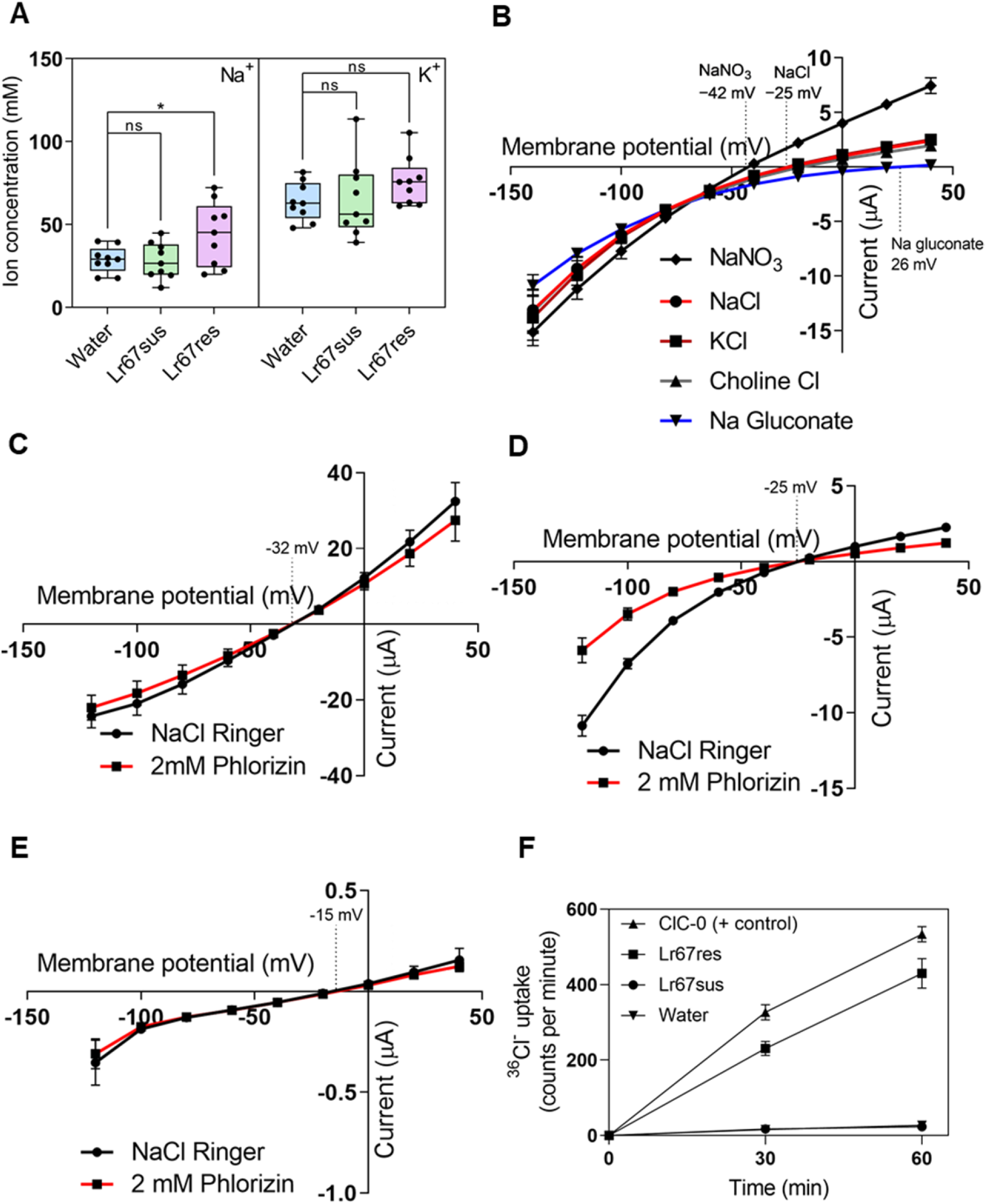
Characterizing the Lr67res gain-of-function in oocytes. **A**, Na^+^ and K^+^ content of oocytes injected with *Lr67sus, Lr67res* or water, bathed in ND96 solution with antibiotics for 48 h post-injection; \**p*<0.05 (t-test comparing *Lr67* alleles with water control); ns, not significant. Current-voltage relationship of **B**, *Lr67res*-injected oocytes perfused with 115 mM Ringer solutions as indicated; **C**, *TmClC-0*; **D**, *Lr67res*; **E**, *Water*-injected oocytes perfused with 115 mM NaCl Ringer ± 30-second 2 mM phlorizin treatment. **F**, Time-course uptake of radiolabeled ^36^Cl^−^ into injected oocytes as indicated, bathed in ND96 without antibiotics at pH 5.5. Boxes in **A** correspond to values within the 25th and 75th percentiles and the horizontal line represents the median; whiskers show max and min values, n=9 pooled oocyte samples. Dotted lines in **B**-**E** indicate reversal potential values. Data points with vertical bars represent the mean ± SE of 4–8 (**B**-**E**) or 9-10 (**F**) oocytes.

The currents detected in *Lr67res* oocytes were compared with those induced by the *Torpedo marmorata* chloride channel (ClC-0) (Jentsch et al., 1990). As found for *Lr67res* oocytes, the *CIC-0*-injected oocytes generated large currents that reversed near −30 mV which is close to the predicted Cl^−^ reversal potential (Weber, 1999) (Figure 3C). Lr67res currents were sensitive to phlorizin (Figure 3D), a known inhibitor of Lr67sus (Moore et al., 2015) and other sugar transporters, whilst the currents in *ClC-0* and control oocytes were unaffected by phlorizin (Figure 3C,E), suggesting the Lr67res protein directly mediates the observed currents. The contribution of anions to the current was further supported by radio-tracer experiments which found that ^36^Cl^−^ uptake by *Lr67res* and *ClC-0-*injected oocytes was comparable and much greater than *Lr67sus* oocytes and water-injected controls (Figure 3F).

### Structure-function analysis of Lr67-like variants based on yeast NaCl sensitivity

To perform structure-function studies of Lr67res, we used the observed accumulation of Na^+^ by *Lr67res* oocytes to develop a phenotypic screen in yeast based on sensitivity to Na^+^. On standard media, the growth of EBY.VW4000 yeast cells (Wieczorke et al., 1999) harboring Lr67res or Lr67sus was similar to the empty-vector controls, but in the presence of 300 mM NaCl, growth of Lr67res yeast was inhibited more than yeast harboring Lr67sus or the empty pDR196 vector (Figure 4A). Enhanced sensitivity to NaCl was also observed in B31 yeast (Bañuelos et al., 1998), unable to efflux Na^+^, harboring Lr67res (Supplemental Figure S2A), and in EBY.VW4000 yeast expressing the *AtHKT1*, known to transport Na^+^ (Uozumi et al., 2000) (Figure 4A). These results are consistent with Lr67res expression in yeast being associated with increased Na^+^ uptake. Lr67res yeast was more sensitive to NaCl, Na_2_SO_4_ and LiCl but not to KCl, K_2_SO_4_ or mannitol (Figure 4A). Lr67res yeast was also sensitive to KI, presumably due to I^−^ anion permeability and toxicity to yeast (Greaves et al., 1928) (Figure 4B). Interestingly, EBY.VW4000 yeast harboring the TmClC-0 chloride channel were also sensitive to NaCl (Figure 4A). Sensitivity to NaCl correlated with *Lr67res* expression since the strong *PMA1* promoter induced sensitivity to NaCl whereas the weaker *ADH* promoter did not (Supplemental Figure S2B). Additionally, N- and C-terminal tags on Lr67res reduced the sensitivity to Na^+^ (Supplemental Figure S2C). Lr67res was unable to rescue the K^+^ uptake-deficient yeast strain CY162 (Ko and Gaber, 1991) (Supplemental Figure S2D) and the K^+^ concentration in Lr67res oocytes was not statistically significantly different from Lr67sus or water-injected oocytes (Figure 3A), indicating that Lr67res is unlikely to facilitate K^+^ transport.

**Figure 4.**
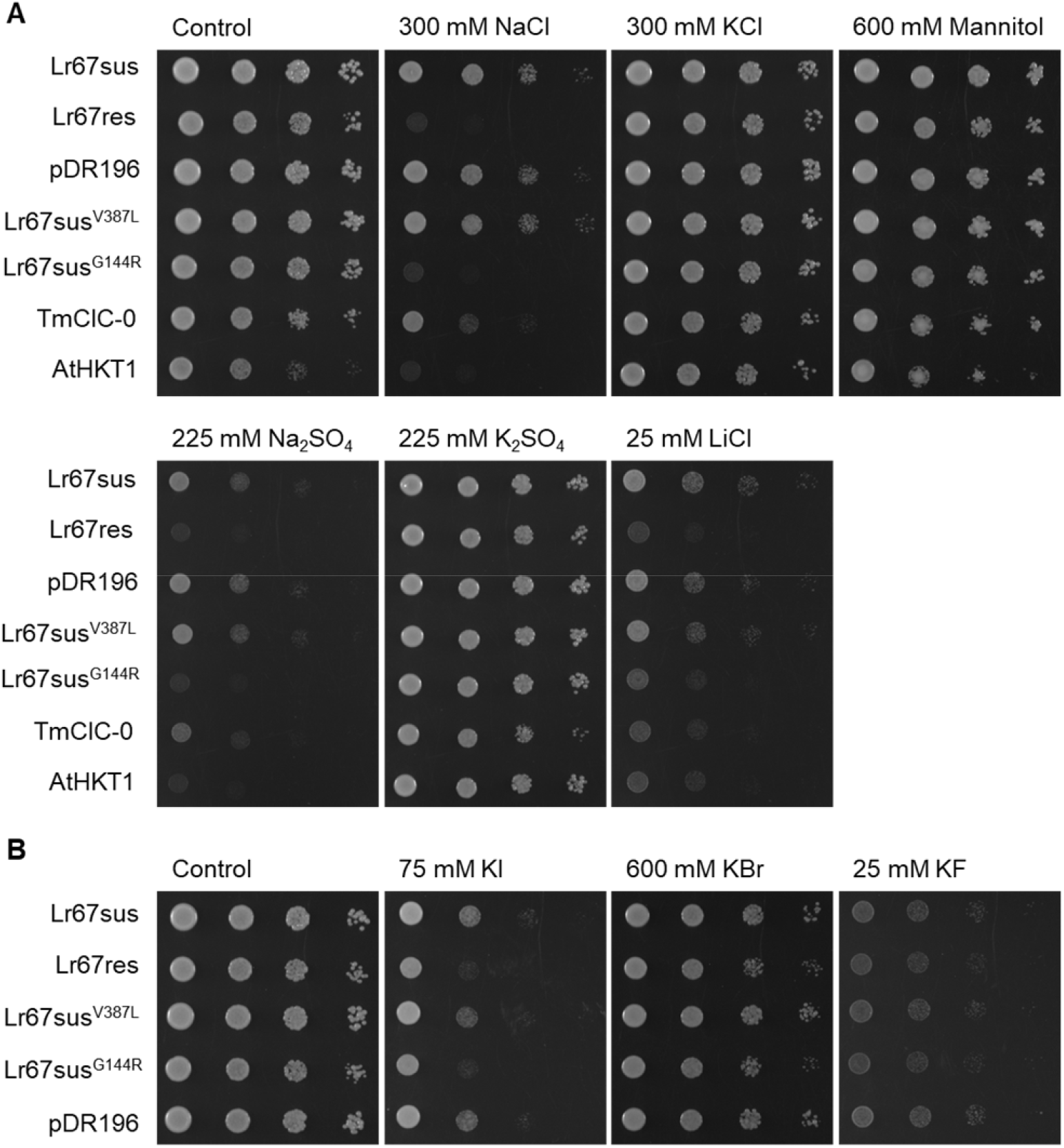
Assessment of Lr67res-induced ion sensitivity in yeast. Decimal dilution series of EBY.VW4000 yeast transformed with *Lr67* alleles, *Lr67* site-directed single mutants, pDR196 empty vector, *Torpedo marmorata* chloride channel, *TmClC-0* and the Arabidopsis high affinity K^+^ transporter, *AtHKT1* grown on media containing different salts to screen for **A**, cation sensitivity or **B**, anion sensitivity. Mannitol was included as an osmotic control. Images are representative of three biological replicates (independently transformed colonies).

### Analysis of Lr67res loss-of-resistance mutants

Four loss-of-resistance (LOR) Lr67res mutants that were identified in wheat (Spielmeyer et al., 2013; Moore et al., 2015) (Lr67res^C75Y^, Lr67res^G208D^ Lr67res^E160K^, Lr67res^G217D^; Figure 5A) cause a loss of Lr67res-mediated resistance and a loss of leaf tip necrosis (Ltn). These LORs were utilized to correlate disease resistance *in planta* with the phenotypes detected in yeast and oocytes. When expressed in yeast these LOR mutants reduced the sensitivity to NaCl (Figure 5B) while maintaining the loss of glucose transport capacity (Figure 5C). Further, when two LOR mutants (*Lr67res*^*C75Y*^, *Lr67res*^*G208D*^) were expressed in oocytes, the large currents typically seen for Lr67res were absent for both mutants (Figure 5D), thus providing a correlation between the observed Lr67res phenotype in oocytes (large inward currents) with disease resistance *in planta*. These results indicate that the gain-of-function phenotype associated with *Lr67res* expression (characterized by enhanced ion fluxes in oocytes and increased NaCl sensitivity in yeast) correlates with Lr67res pathogen resistance in wheat. Co-expression of *Lr67sus* in yeast with either *Lr67res, Lr67res*^C75Y^ or *Lr67res*^G208D^ resulted in ∼50% lower glucose uptake compared to *Lr67sus* alone (Figure 5E) and co-expression of *Lr67res* with *Lr67sus* in yeast maintained NaCl sensitivity (Figure 5F). Lr67sus glucose uptake was unaffected by co-transformation with an empty vector, most likely because the empty vector did not produce any protein (Figure 5E).

**Figure 5.**
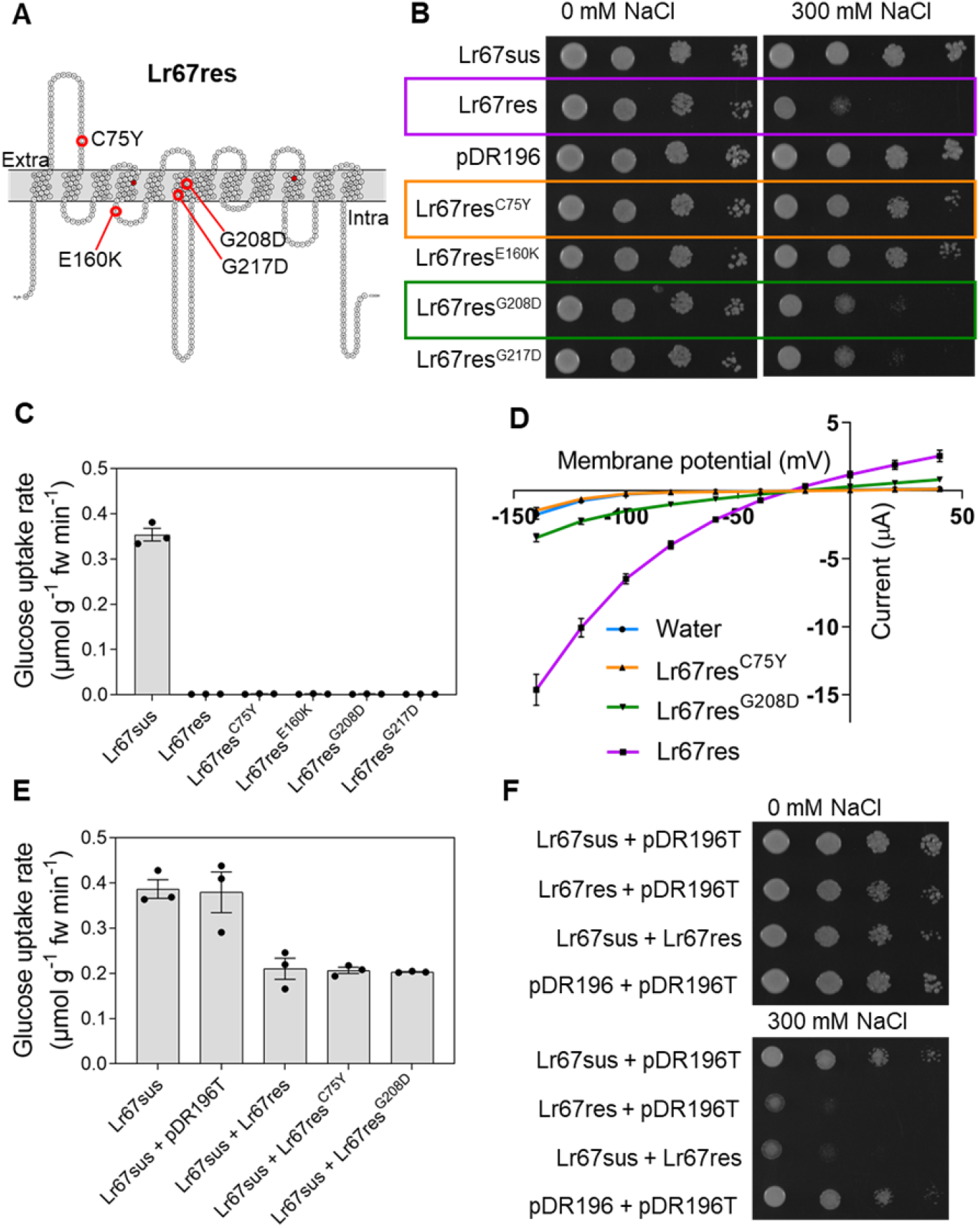
Lr67res loss-of-resistance (LOR) mutants do not transport glucose and lack the Lr67res gain-of-function. **A**, Predicted transmembrane topology plot highlighting four single amino acid changes that cause a loss-of-resistance phenotype in *Lr67res* wheat (Moore et al., 2015), extracellular and intracellular domains are indicated along with G144R and V387L. **B**, Decimal dilution series of EBY.VW4000 yeast transformed with *Lr67sus, Lr67res*, pDR196 empty vector or *Lr67* loss-of-resistance mutants in **A** grown on media supplemented with NaCl as indicated. **C**, Four-minute [^14^C]-glucose uptake into EBY.VW4000 yeast transformed with *Lr67sus, Lr67res* or each mutant in **A. D**, Current-voltage relationship of *X. laevis* oocytes perfused with 115 mM NaCl Ringer solution under voltage-clamped conditions, injected with water, *Lr67res, Lr67res*^*C75Y*^ or *Lr67res*^*G208D*^. **E**, Four-minute [^14^C]-glucose uptake of yeast transformed with *Lr67sus* alone or *Lr67sus* co-transformed with pDR196T empty vector, *Lr67res* or LOR mutants as indicated. **F**, Decimal dilution series of EBY.VW4000 yeast co-transformed with *Lr67* alleles and pDR196/pDR196T empty vectors. Images (**B, F)** are representative of three biological replicates (independently transformed yeast colonies). Columns with vertical bars (**C, E)** represent mean ± SE of three biological replicates (independently transformed yeast colonies). Data points with vertical bars (**D**) represent the mean ± SE of 5-8 oocytes.

### The enhanced NaCl sensitivity in yeast can be induced by other members of the STP13 family and residues other than arginine are capable of mirroring the Lr67res phenotype

Since the first two *Lr67* alleles identified in wheat (*Lr67res* and *Lr67sus*) differ by two residues (G144R and V387L), we tested whether both changes are required to induce NaCl sensitivity in yeast. We established that the single G144R mutation was sufficient to induce NaCl sensitivity in yeast but the single V387L mutation could not (Figure 4A). This is consistent with the more recent identification of another naturally occurring Lr67 variant that also confers resistance but that contains the G144R mutation only. When this G144R allele was crossed into the wheat cultivar Avocet S, which is highly susceptible to stripe rust, it conferred the same partial resistance to wheat stripe rust as conferred by the *Lr67res* allele with both G144R and V387L mutations (Figure 6A). Other amino acid substitution variants tested in yeast (Lr67sus^G144C^ and Lr67sus^G144D^) also exhibited increased NaCl sensitivity, indicating that the change from glycine to arginine is not critical for this gain-of-function phenotype (Figure 6B). Since the Lr67sus^G144A^ variant did not confer NaCl sensitivity, amino acids of a particular size or charge may be required to induce this phenotype.

**Figure 6.**
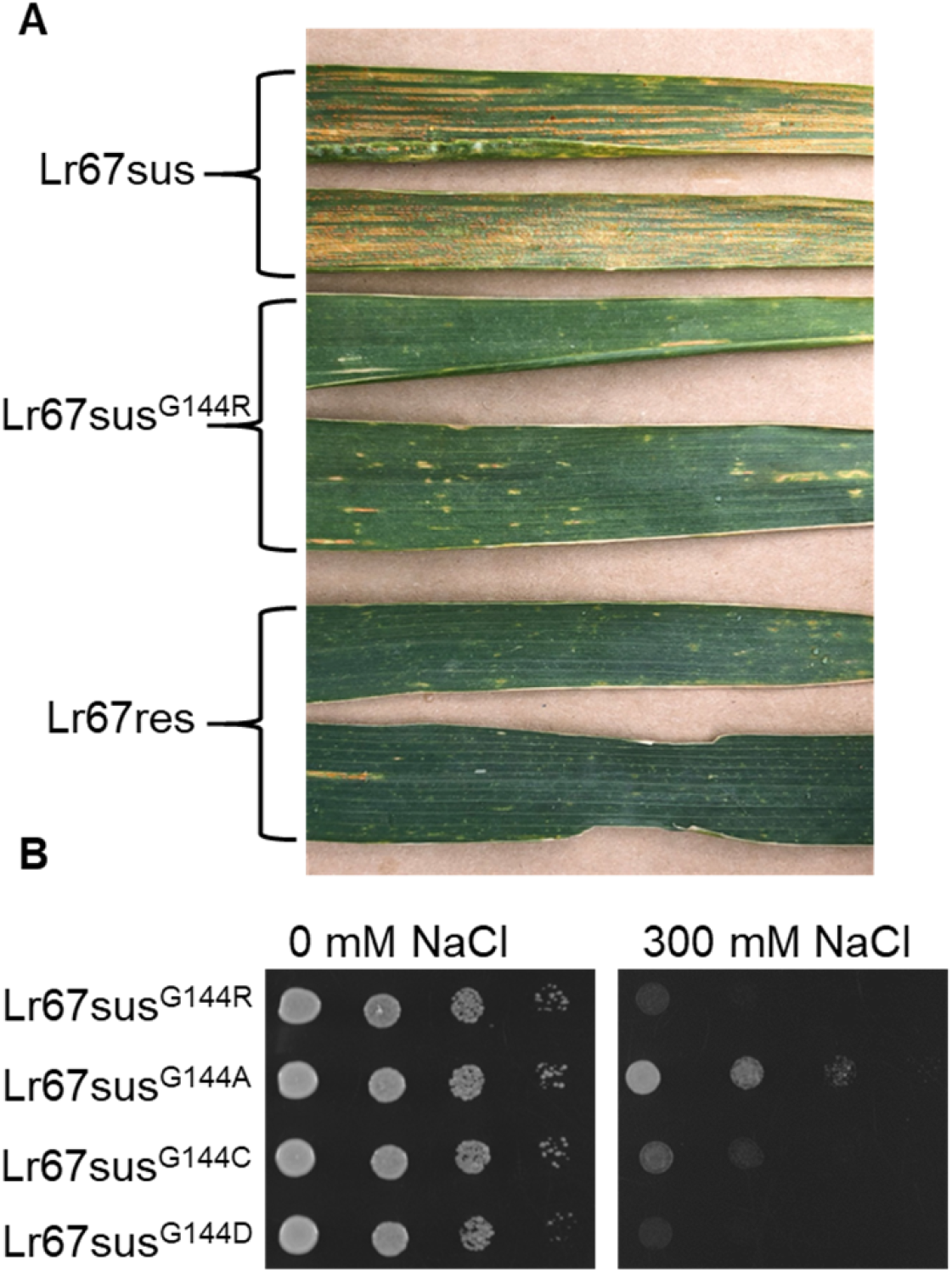
Disease resistance conferred by the G144R mutation in wheat and yeast NaCl sensitivity of site-directed mutants. **A**, Flag leaves of field-grown cv. Avocet S wheat carrying *Lr67sus, Lr67sus*^*G144R*^ or *Lr67res*, infected with *P. striiformis* f. sp. *tritici*. **B**, Decimal dilution series of EBY.VW4000 yeast transformed with *Lr67sus*^*G144R*^, *Lr67sus*^*G144A*^, *Lr67sus*^*G144C*^, *Lr67sus*^*G144D*^ site-directed mutants grown on media supplemented with NaCl as indicated, image is representative of three biological replicates (independently transformed yeast colonies).

Since the G144 residue is highly conserved in plant sugar transporters it could provide a novel strategy for introducing disease resistance into other species. Therefore, we introduced orthologous mutations into transporters from other species, expressed them in yeast, and tested whether they conferred the same sensitivity to NaCl. Introducing the single G144R mutation into the barley *HvSTP13* gene was sufficient to confer NaCl sensitivity to yeast and no additional sensitivity was evident when G144R was combined with the V387L mutation (Supplemental Figure S3). Furthermore, the *Arabidopsis* AtSTP13^G145R^ protein and, to a lesser extent, the sorghum SbSTP13^G144R^ protein also exhibited increased NaCl sensitivity in yeast (Figure 7). These same variants were unable to complement yeast growth on glucose media suggesting they too lack the hexose transport function as demonstrated previously for Lr67res (Moore et al., 2015) and HvSTP13^G144R;^ (Milne et al., 2019) (Figure 7). By contrast, similar mutations introduced to the structurally-resolved bacterial xylose-proton symporter (Sun et al., 2012) (XylE^G137R^) and variants of the human glucose facilitator (Wang et al., 2005; Deng et al., 2014) (GLUT1^G130R^ and GLUT1^G130S^) did not enhance NaCl sensitivity (Figure 7). These results suggest that generation of the NaCl-sensitive phenotype in yeast may only be possible in closely related plant transporters within the STP family. In this context, the synthetic barley, Arabidopsis, and sorghum variants that conferred NaCl sensitivity to yeast were between 98.8% and 81.4% identical to Lr67, whereas the bacterial XylE^G137R^ and human GLUT1^G130R^ shared less identity (∼27% identical). More specifically, an extracellular disulphide bond in the lid domain between Cys77 and Cys449 identified in the AtSTP10 crystal structure (Paulsen et al., 2019) may be required for the Lr67res gain-of-function. The equivalent C75Y LOR mutation of Lr67res abolished resistance in wheat (Spielmeyer et al., 2013; Moore et al., 2015) and disrupted the gain-of-function in yeast and oocytes (Figure 5B,D). Moreover, the corresponding amino acid is not conserved in XylE or GLUT1 which may explain why XylE^G137R^ and GLUT1^G130R^ did not phenocopy the yeast NaCl sensitivity.

**Figure 7.**
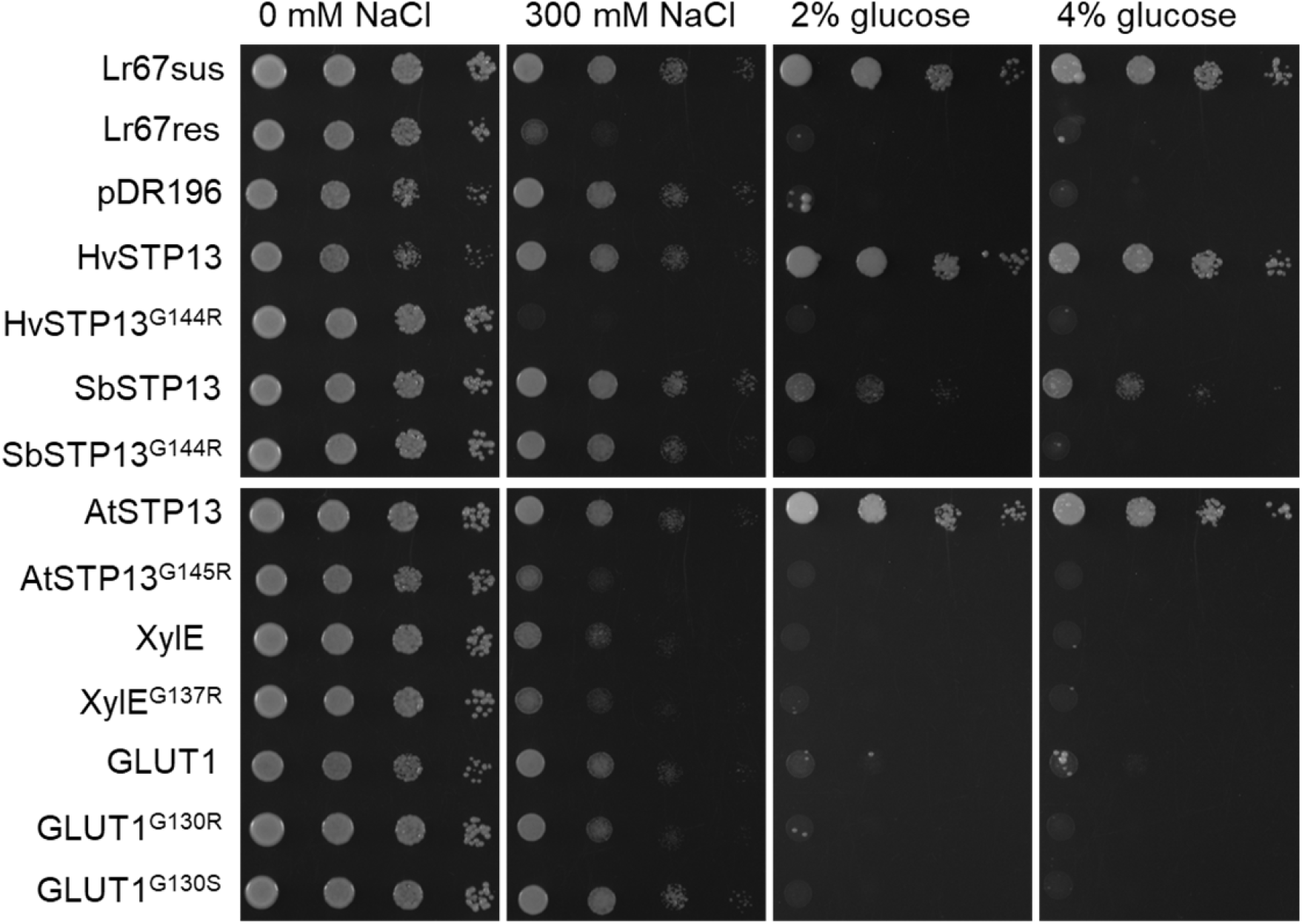
NaCl sensitivity and glucose uptake capability of yeast transformed with *Lr67*-related hexose transporters and site-directed mutants. Decimal dilution series of EBY.VW4000 yeast transformed with the wheat *Lr67sus* and *Lr67res*, empty vector pDR196, barley *HvSTP13* and *HvSTP13*^*G144R*^, sorghum *SbSTP13* and *SbSTP13*^*G144R*^, Arabidopsis *AtSTP13* and *AtSTP13*^*G154R*^, *E. coli XylE* and *XylE*^*G137R*^, human *GLUT1, GLUT1*^*G130R*^ and *GLUT1*^*G130S*^. Media was supplemented with NaCl as indicated, or maltose was substituted for glucose as the carbon source. Images are representative of three biological replicates (independently transformed yeast colonies).

### Extending observations from heterologous systems to plants

To determine whether salt treatments could induce distinct phenotypes in wheat plants expressing different alleles of *Lr67*, as they did in yeast, near isogenic cv. Thatcher wheat lines carrying either the *Lr67sus* or *Lr67res* alleles (Thatcher and Thatcher+Lr67res respectively) were subjected to NaCl treatment during the adult stage of development. According to our observations, Thatcher+Lr67res plants do not normally display partial disease resistance or the leaf tip necrosis (Ltn) phenotype when grown in glasshouses or growth cabinets, rather, these phenotypes only occur in field-grown adult plants. However, we were able to induce Ltn in the flag leaves of cabinet-grown Thatcher+Lr67res adult plants with NaCl treatments commencing at anthesis (25 mM followed by 50 mM NaCl). Induction of Ltn did not occur in Thatcher plants, or in plants of both genotypes that did not receive salt treatment (Figure 8A; Supplemental Figure S4). Leaf tip necrosis was also evident on the penultimate leaves and older leaves of NaCl-treated Thatcher+Lr67res (not shown) indicating, as for yeast and oocytes, an Lr67res-dependent ion-responsive phenotype is transferrable to plants.

**Figure 8.**
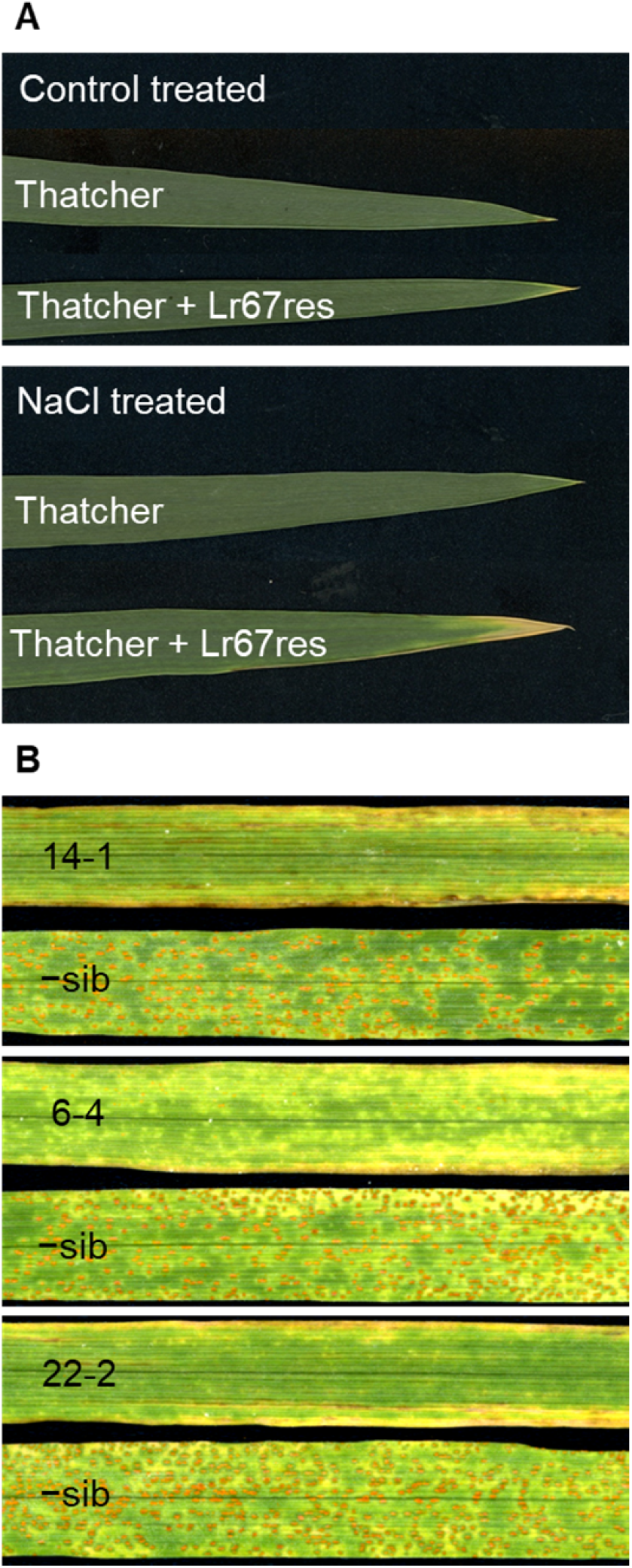
NaCl induced leaf tip necrosis; rust resistance phenotype of barley transformed with *HvSTP13*^*G144R,V387L*^. **A**, Flag leaves collected from the main tiller of representative wheat cv. Thatcher or Thatcher+Lr67res plants treated with half-strength Hoaglands solution ± NaCl. A 25 mM NaCl treatment was applied at anthesis, followed by 50 mM NaCl treatment 6 days later and leaves were sampled 5 days after the second treatment. Images are representative of 12 biological replicates and two independent experiments. Additional replicates are shown in Supplemental Figure S4. **B**, Barley leaves from three independent transgenic events inoculated with *P. hordei*. Images represent leaves of barley plants carrying the *HvSTP13*^*G144R,V387L*^ transgene, or corresponding −sib lacking the transgene at 8 days post-inoculation.

To further translate the observations made with the synthetic *HvSTP13* variants in yeast to whole plants, a construct was prepared containing the full genomic sequence of the modified *HvSTP13* with both mutations (*HvSTP13*^*G144R,V387L*^) along with 1503 bp of its native promoter and a 1660 bp 3’ UTR. The construct was transformed into barley cv. Golden Promise and T1 plants underwent rust disease screening. Multiple independent transgenic events containing the *HvSTP13*^*G144R,V387L*^ transgene exhibited partial disease resistance to barley leaf rust when inoculated with *Puccinia hordei*, with three independent events presented, whereas their non-transgenic sibling lines (−sibs) remained susceptible (Figure 8B). Microscopic analysis distinguished between strong and intermediate resistance phenotypes at 8 dpi (Figure 9A-H) which became visible macroscopically at 13 dpi (Figure 9I). Hyphal development and sporulation were evident in −sibs lacking *HvSTP13*^*G144R,V387L*^, some hyphal development in the absence of sporulation was observed in intermediately resistant transgenic events, whereas only minimal hyphal development was evident in the strongly resistant transgenic events. If only the single G144R change (or mutation to other amino acids) is required as indicated by yeast results (Figure 7A) and demonstrated in wheat (Figure 6A), this will simplify the introduction of Lr67-like resistance to other crops in the future.

**Figure 9.**
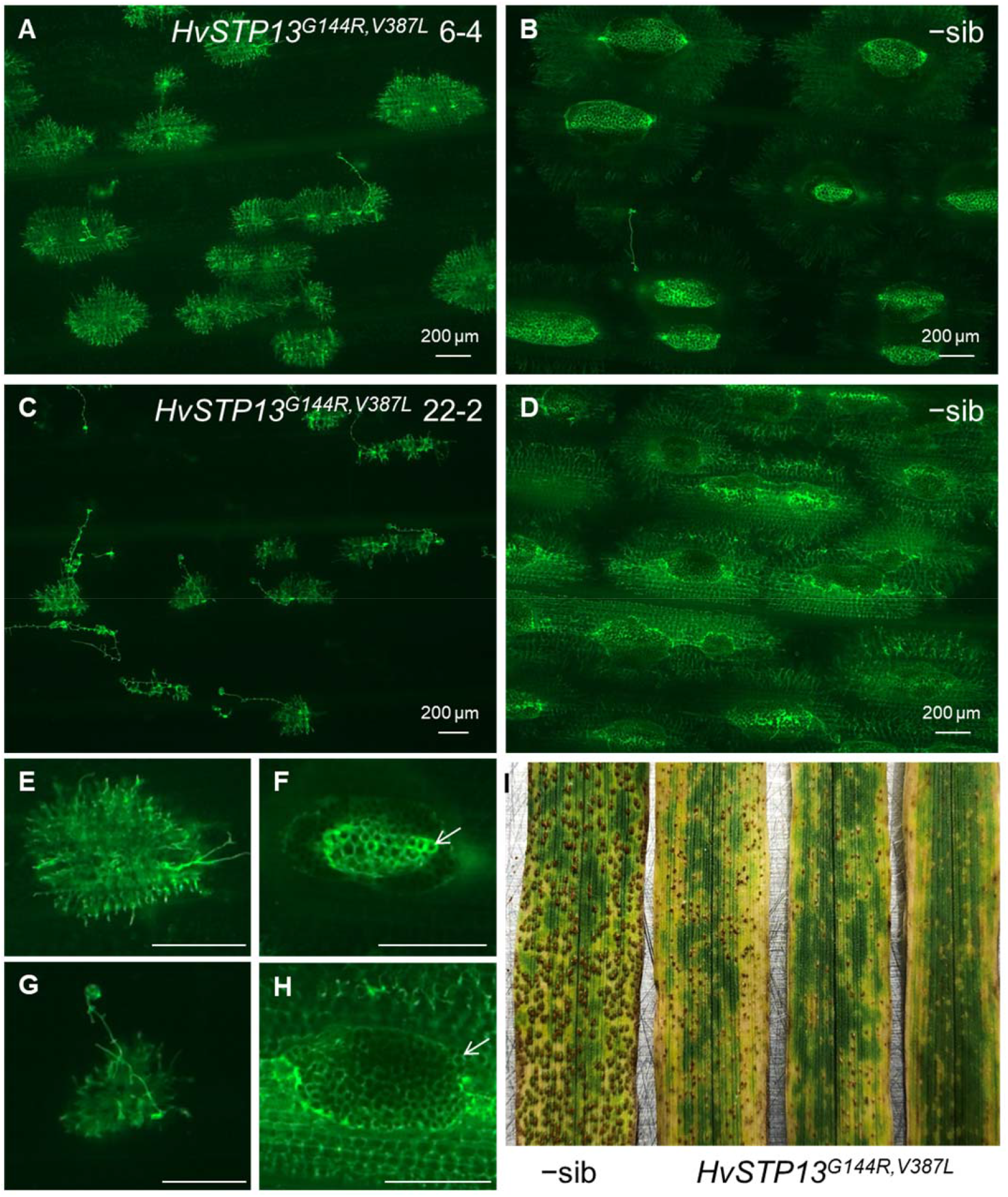
Fluorescence micrographs of WGA-FITC stained barley cv. Golden Promise leaves taken 8-days post-inoculation with *P. hordei*. Representative images for plants exhibiting **A** intermediate and **C** strong resistance phenotypes harvested from transgenic lines expressing *HvSTP13*^*G144R,V387L*^. **B**,**D**, Susceptible corresponding segregant, −sibs, lacking the *HvSTP13*^*G144R,V387L*^ transgene, exhibiting extensive fungal colonization and sporulation. Higher magnification images **E**-**H** of infection sites corresponding to **A**-**D** respectively depicting presence (**F**,**H**; white arrows) or absence **E**,**G** of sporulation. Minimal and moderate hyphal development was observed in **C** and **A** respectively (magnified in **G** and **E**) in the absence of sporulation. All scale bars = 200 μm. **I**, Infected leaves at 13 dpi from −sib and independent *HvSTP13*^*G144R,V387L*^ transgenic events exhibiting various levels of sporulation on transgenic plants.

## DISCUSSION

### Considering the feasibility of a sugar transporter exhibiting ion channel-like properties

This study demonstrates that the Lr67res variant confers a gain-of-function phenotype that changes the permeability of the plasma membrane to small ions. The observed shifts in oocyte reversal potential (Figure 2B) and the close parallels with observed properties of the well-characterized chloride channel TmClC-0 in both yeast and oocyte systems (Figures 3C, 4A) provide evidence that anion fluxes, particularly chloride, may account for the Lr67res-dependent currents in oocytes. While it is impossible to eliminate the involvement of endogenous oocyte transporters in mediating this flux, the inhibition of current by the sugar transporter inhibitor phlorizin (Figure 3D) strengthens the argument that the observed currents are indeed Lr67res-mediated. Ion sensitivity in the form of NaCl-induced Ltn in Thatcher+Lr67res wheat further supports this argument (Figure 8A). We closely examined the prospect of observed oocyte currents being a product of uncoupled flux of H^+^ through Lr67res, however the hallmarks of H^+^ flux that were observed for the functional Lr67sus symporter were absent for Lr67res (Figure 2E,F). Thus, we concluded that protons are unlikely to be responsible for the observed current, and this is discussed in more detail below.

It is conceivable that mutation of the highly conserved G144 residue of Lr67 does indeed confer a gain-of-function and simultaneous loss of sugar transport function given observations from previous studies. Most significant of these are recent reports of a transient Cl^−^ binding site identified in the crystal structure of an inward-open conformation of the closely related AtSTP10 (Bavnhøj et al., 2021) and more phylogenetically-distant glucose transporter GLUT1 (Custódio et al., 2021). We suggest that the G144R mutation of Lr67res may enable the transit of this bound Cl^−^ through the transporter, giving rise to the novel gain-of-function reported in our study. Additionally, the Lr67res G144R mutation is found one helical turn from conserved arginine and aspartate residues of the structurally resolved XylE (Arg133, Asp27) and AtSTP10 (Arg142, Asp42) that constitute the proton binding site of each transporter (Sun et al., 2012; Paulsen et al., 2019). The protonation/deprotonation of this pair is thought to drive the conformational change that drives hexose transport (Wisedchaisri et al., 2014). The close proximity of G144R to these critical residues within the predicted 3D structure of Lr67res supports our observations of Lr67res having a gain-of-function. There are also precedents for single residue mutations, or a 4-residue deletion in the case of hGLUT1 (Weber et al., 2008), in transporters beyond the SP family substantially altering protein function. For instance, single residue mutations can uncouple the transport of substrates from their co-transported ions (Borre and Kanner, 2004; Martial et al., 2006), or induce novel channel-like properties that allow the passage of previously excluded ions (Meredith, 2004; Bruce et al., 2005; Weber et al., 2008; Qin and Boron, 2013). Also of particular relevance is that a single amino acid mutation in the proton-pumping bacteriorhodopsin converts it from a H^+^ pump to a Cl^−^ pump (Sasaki et al., 1995).

### Neither protons or a loss of glucose transport are unlikely to underpin Lr67res associated ion fluxes

As Lr67sus is a proton-coupled hexose transporter, the possibility of a simple uncoupling of glucose and proton transport for Lr67res was closely examined. We showed that glucose uptake by *Lr67sus*-injected oocytes was associated with glucose-induced inward currents, glucose and pH-dependent shifts in reversal potential and cytosolic acidification (Figures 1A, 2B) confirming at proton fluxes were detectable in the *X. laevis* expression system. The lack of glucose uptake by *Lr67res*-injected oocytes (Figure 1B) agrees with previous observations in yeast (Moore et al., 2015), and while the current magnitude increased in acidic solutions, there was no concurrent shift in reversal potential (Figure 2E) and no differences in the rate of cytosolic acidification between *Lr67sus, Lr67res* or control oocytes when external pH was acidified (Figure 2A, Supplemental Figure S1). Thus, protons were unlikely to be a major contributor to the large observed current in *Lr67res* oocytes. Although *Lr67res* yeast cells were sensitive to NaCl and *Lr67res-*injected oocytes accumulated more Na^+^, electrophysiological studies indicated that other cations were not a major contributor to the currents either – the minor changes in current magnitude when Na^+^ was substituted for K^+^ or choline^+^ and by the shifts in reversal potential associated with changing external NaCl concentration were more consistent with anion permeability. Furthermore, the currents measured in *Lr67res* oocytes contrasted with those induced by the expression of two known Na^+^ transporters, TaHKT1;5-A and AtHKT1. These proteins caused a positive shift in reversal potential as external Na^+^ was increased (Uozumi et al., 2000; Munns et al., 2012), whereas a negative shift was observed in *Lr67res* oocytes. To reconcile the Lr67res-induced NaCl sensitivity of yeast with the large anion fluxes in oocytes, we propose that under high external NaCl in yeast, the uptake of Cl^−^ by *Lr67res* yeast triggers the uptake of Na^+^ to maintain electroneutrality, with accumulation of Na^+^ continuing until it becomes toxic. This is consistent with NaCl but not KCl inducing toxicity and is supported by the observation that *TmClC-0*-expressing yeast shows the same NaCl sensitivity (Figure 4A), albeit to a slightly lesser extent than *Lr67res* yeast which may be due to differing expression levels. Since elevated Na^+^ concentration in oocytes was observed (Figure 3A), a similar uptake of Na^+^ to maintain electroneutrality may also occur.

Our experimental evidence does not support the model of a loss of glucose transport function by Lr67res underpinning disease resistance. Firstly, mutations of Lr67res that caused a loss of resistance *in planta* (C75Y, E160K, G208D, G217D) (Spielmeyer et al., 2013; Moore et al., 2015) did not restore glucose transport by Lr67res *in vitro* (Figure 5A, C). Secondly, the reduction in yeast glucose uptake observed when *Lr67sus* was co-expressed with an LOR mutant resembled that when *Lr67sus* was co-expressed with *Lr67res* (Figure 5E), Together, these results indicate that *in planta* resistance is unlikely to be caused by a dominant-negative interference of glucose uptake, as proposed previously (Moore et al., 2015). Further, the reduction in Lr67res oocyte currents by LOR mutations (Figure 5D) gives further support to correlate the Lr67res gain-of-function with resistance.

### Prospects for engineering novel disease resistance with STP13 mutants

The development of an NaCl sensitivity screen in yeast, which we predict relies on the movement of toxic Na^+^ into yeast cells to maintain electroneutrality under Cl^−^ influx conditions, enabled rapid detection of an Lr67res phenotype without the requirement of specialized electrophysiology equipment. This assay can be adopted as a pre-screening tool to test for an NaCl sensitivity phenotype of synthetic STP13 variants prior to producing transgenic plants, as a proxy for disease resistance. The confirmation that the G144R substitution is the sole requirement for conferral of Lr67res-mediated disease resistance in wheat (Figure 6A) and barley (Skoppek et al., 2022) (independent to this study), and the high conservation of STP13 across plant species makes it a valuable, uncomplicated target for gene editing. Further, the utility of this point mutation represents a valuable target for future gene editing studies. The G144R substitution is rare in wheat genotypes and was not identified in diverse accessions of barley (Milne et al., 2019). If this is also the case in other crops, mutagenesis would be a logical step to develop novel sources of non-transgenic resistance. However, the G144R change may not be readily achieved via sodium azide or EMS chemical mutagenesis, therefore it would be worthwhile mining mutant populations for induced variants such as G144D, which lends itself to being induced chemically into the native sequence of STP13 in several monocot species, and also confers NaCl sensitivity in yeast (Figure 6B). Notwithstanding the issues of pleiotropy in our study, a significant finding is the proof-of-concept that an Lr67 orthologue can be modified and stably transformed in its native species to confer disease resistance (Figures 8B, 9) and the V387L change is superfluous for disease resistance (Skoppek et al., 2022).

### Lr67res and the link between anion fluxes and plant immunity

Based on the data presented, we propose that enhanced ion fluxes are likely to be a biologically relevant factor for *in planta* multi-pathogen disease resistance for two reasons. Firstly, the yeast ion sensitivity phenotypes (or lack of sensitivity in the case of LOR mutants) corresponded in disease resistance phenotypes when tested in both wheat and barley plants. Secondly, the induction of Ltn by NaCl treatment, a phenotype linked with disease resistance phenotypes for the multi-pathogen resistance genes *Lr67, Lr34* and *Lr46* (Singh et al., 1998; Krattinger et al., 2009; Moore et al., 2015), in growth conditions not otherwise conducive to the development of both Ltn and partial disease resistance, suggests that both may be ion-inducible. In the context of disease resistance, anion fluxes, and in particular chloride, have been implicated in immune signaling (Liu et al., 2019), with evidence accumulating of chloride channels and transporters being both positive (Han et al., 2019) and negative (Guo et al., 2014) regulators of PAMP- and effector-triggered immunity. Additionally, ABA, which is the transport substrate of the other characterized multi-pathogen resistance gene, Lr34 (Krattinger et al., 2019), triggers channel mediated Cl^−^ fluxes (Roelfsema et al., 2004), and induces leaf senescence (Mao et al., 2017). Occurrence of Ltn in plants carrying an *APR* gene in the absence of pathogen infection indicates that physiological changes leading to resistance are not necessarily reliant on pathogen induction. Instead, we propose that Lr67res-altered ion fluxes *in planta* underpin both multi-pathogen resistance and the observed Ltn induction, and future experiments will work towards characterizing whether these phenotypic responses are due to localized perturbations in ion distribution or more systemic ion-induced abiotic stress signaling. In terms of general functionality of multi-pathogen resistance proteins in wheat, Lr67 appears to possess a distinct molecular function to *Lr34*, since Lr34 did not induce the same sensitivity to NaCl (Supplemental Figure S2), nor did it induce large currents in oocytes (Figure 2D), which reconciles with involvement of Lr34 in ABA transport (Krattinger et al., 2019). In contrast to the classical immune response involving gene-for-gene recognition which has recently been linked to calcium ion permeability culminating in the hypersensitive response and cell death (Bi et al., 2021; Jacob et al., 2021), the partial multi-pathogen resistance and durability of both Lr67res and Lr34res could instead rely on changes to ion permeability of the host cell’s plasma membrane, rendering the host tissues less conducive to pathogen growth and virulence, in the absence of the hypersensitive response, through an as yet unidentified mechanism.

In summary, we have identified and characterized a novel gain-of-function that the Lr67res protein displays over the Lr67sus protein, which leads to novel ion fluxes and ion sensitivity in different biological systems. This study provides the first detailed characterization of the Lr67res gain-of-function and presents the significant finding that this Lr67 gain-of-function can be phenocopied in yeast using STP13 transporters from other species and *in vivo* in barley. This signifies STP13, and potentially other related STPs, as promising targets for chemical mutagenesis or gene editing to improve the disease resistance of crops in the future.

## MATERIALS AND METHODS

### Constructs for yeast and oocyte expression

The full-length coding sequences of *Lr67* and *Lr34* alleles were PCR amplified and cloned into the pENTR1A entry vector (Life Technologies, Mulgrave, VIC, Australia) using specified primers and restriction sites (Supplemental Table S1). Site-directed mutagenesis was used to introduce C75Y, E160K, G208D and G217D mutations into *Lr67res* using primers in Supplemental Table S1, designed by the QuikChange webpage – https://www.genomics.agilent.com/primerDesignProgram.jsp using the protocol as described for *HvSTP13* (Milne et al., 2019). Each gene was recombined into the destination oocyte expression vector, pGEMHE-DEST (Shelden et al., 2009), using LR Clonase Recombinase (Life Technologies) according to the manufacturer’s protocol. Gene sequences were confirmed by Sanger sequencing (AGRF, Westmead, NSW, Aust). Thereafter, plasmids were linearized with the NheI restriction enzyme (NEB, Ipswich, MA, USA) to transcribe complimentary RNA (cRNA) driven by the T7 promoter using the Ambion mMessage mMachine kit (Life Technologies) according to the manufacturer’s protocol.

For heterologous expression in yeast, *Lr67* and *Lr34* alleles were sub-cloned from pENTR1A into either the pDR195 or pDR196 yeast vector (Rentsch et al., 1995) using EcoRI and XhoI, or BamHI and XhoI restriction sites respectively. *AtSTP13* (NotI, XhoI), *SbSTP13* (EcoRI, XhoI), *XylE* (EcoRI, XhoI), *AtHKT1* (NotI, BamHI) and *Torpedo marmorata chloride channel* (*ClC-0*; SpeI, XhoI) cDNAs were synthesized by Integrated DNA Technologies (Singapore Science Park II, Singapore) with flanking restriction sites as indicated. *GLUT1* (NotI, BamHI) was synthesized by GeneArt (ThermoFisher, Scoresby, VIC, Australia). Constructs are detailed in Supplemental Table S2. Site-directed mutants were produced after cloning into pENTR1A using the primers in Supplemental Table S1 and protocol above, and then sub-cloned using flanking restriction sites into pDR195 or pDR196. FLAG-tagged *Lr67* alleles in p426ADH1 as described (Moore et al., 2015) were sub-cloned into pDR196 using EcoRI and SalI restriction sites. Yeast expression constructs were transformed into yeast as described (Dohmen et al., 1991), at least three colonies were selected as independent transformation events (biological replicates). Colonies were cultured in synthetic dropout medium lacking uracil (SDura^−^; 6.72 g/L yeast nitrogen base with ammonium sulfate, 0.96 g/L yeast synthetic dropout medium without uracil, 2% w/v maltose), plasmids were rescued from yeast as described using the QIAprep Miniprep Kit (QIAGEN, Chadstone Centre, VIC, Australia – User developed protocol: Isolation of plasmid DNA from yeast), transformed to *E. coli* and sequences were confirmed by Sanger sequencing (AGRF). Yeast strains harboring constructs of *Lr67* alleles driven by the yeast *ADH1* promoter in p426ADH1 were used as described (Moore et al., 2015).

### Expression in oocytes, radiolabeled uptake and two-electrode voltage-clamp (TEVC) experiments

Stage V–VI oocytes were selected and injected with 46 ng of cRNA or an equal volume (46 nL) of RNA-free water and incubated in Frog Ringers solution supplemented with antibiotics (96 mM NaCl, 2 mM KCl, 1 mM MgCl_2_, 0.6 mM CaCl_2_, 5% v/v horse serum, 100 units/mL penicillin, 1 mg/mL streptomycin, 0.5 mg/mL tetracycline, HEPES-NaOH pH 7.6) at 18°C prior to experimentation. ^36^Cl^−^ uptake was performed two days post-injection (dpi) and oocytes were incubated in Frog Ringers solution at pH 5.5 without antibiotics. Water, *Lr67sus, Lr67res* and *TmClC-0* injected oocytes were incubated at room temperature in Frog Ringers solution containing H[^36^Cl^−^]. Oocytes were washed three times by pipetting into 2 mL of ice-cold Frog Ringers solution lacking ^36^Cl^−^. [^14^C]-glucose uptakes were performed identically to ^36^Cl^−^ uptakes but *Lr67sus, Lr67res* and water-injected oocytes at 2 dpi were incubated for one hour in 115 mM NaCl MES-Tris pH 5 Ringer solution containing 11.4 μM [^14^C]-glucose and washes were performed using ice cold 115 mM NaCl MES-Tris pH 5 Ringer solution supplemented with 1 mM glucose. Each oocyte was placed in a separate scintillation vial, dissolved in 0.1% nitric acid and 4 mL Optima Gold XR scintillant (Perkin Elmer, Glen Waverley, VIC, Australia) was added prior to counting for two minutes using a Liquid Scintillation Counter (LS6500; Beckman Coulter, Lane Cove, NSW, Australia).

Glucose uptake of Lr67sus was performed using 2 dpi oocytes that were placed in a recording bath and perfused with a modified Na-Ringer solution (MES-Tris Ringer; 115 mM NaCl, 2 mM KCl, 1.8 mM CaCl_2_, 1 mM MgCl_2_, 5 mM MES-Tris at appropriate pH) (Sivitz et al., 2007). Recording pipettes, 3 M KCl filled and ∼1 megaohm resistance, measured currents using the two-electrode voltage-clamp technique as described (Sivitz et al., 2005). Pulses were applied for 203 ms at voltages from –140 mV to 40 mV in 20 mV increments. Steady-state currents are presented as the mean current between 150 and 200 ms following the onset of voltage pulses. Substrate-dependent currents were obtained by subtracting an averaged background current before and after the provision of substrate. Solution osmolarities were adjusted using mannitol to 240–260 mOsmol kg^−1^ using a Wescor vapor pressure osmometer. Due to the magnitude of currents obtained in Lr67res oocytes and deterioration of oocytes at 2 dpi, measurements were carried out 1 dpi and presented as IV curves. Glucose induced currents by Lr67sus were detectable at this time point and were observed to increase between 1 and 3 dpi. Ringer solution was as described unless indicated. For phlorizin inhibition experiments, phlorizin was first dissolved in DMSO and added to MES-Tris Ringer at pH 5, to a final concentration of 2 mM phlorizin and 1% DMSO. The control solution lacking phlorizin also contained the same concentration of DMSO and was deemed to not have an impact on Lr67res currents.

### Quantification of oocyte ion content

Water, *Lr67sus* and *Lr67res*-injected oocytes incubated at 18°C in Frog Ringers with antibiotics were sampled 48 hours after injection. Two oocytes were pooled per sample and oocytes from two frogs were sampled. Oocytes were washed twice with Milli-Q water which was removed by pipetting. Oocytes were frozen at –20°C, thawed and digested in 100 μL 0.1 M nitric acid at 42°C for 2 h before adjusting total volume to 1 mL with Milli-Q water. Debris was removed by centrifugation and the clarified supernatant was diluted for measurement on an Atomic Absorption Spectrometer (AA-7000F; Shimadzu, North Plympton, SA, Aust). Standard curves were prepared with commercial standards for Na^+^ and K^+^ (Sigma-Aldrich, North Ryde, NSW, Aust) and clarified supernatants were diluted with Milli-Q water accordingly to fit the range of each standard curve. Ion concentration was calculated using 1 μL as the oocyte volume (Kelly et al., 1995).

### Measurement of oocyte cytosolic pH and membrane potential

Cytosolic pH changes were monitored using a proton sensitive microelectrode whilst membrane potential changes were monitored using a voltage electrode (electrodes were prepared as described; Bose et al., 2013). Each oocyte was placed in a recording bath submerged in a modified Na-Ringer solution at pH 7.6 or 4.5 (± 1 mM glucose) that was circulated by a peristaltic pump for ∼5 minutes after solution change before the flow was stopped. Membrane potential and pH measurements were recorded constantly for approximately 60 minutes per oocyte using the MIFE™ (University of Tasmania, Hobart, Australia) system.

### Yeast plate and uptake assays

EBY.VW4000 yeast (Wieczorke et al., 1999) (a *hxt-null* line incapable of hexose uptake) and B31 yeast (Bañuelos et al., 1998) (incapable of Na^+^ efflux) were transformed using the PEG1000 protocol (Dohmen et al., 1991). Transformed yeasts were cultured in liquid SDura^−^ medium to early logarithmic phase (OD_600_ of 0.8 to 1.0) and re-suspended to an OD_600_ of 0.8. A decimal dilution series of yeast suspension (3 μL) was spotted onto solid (15 g/L agar) SDura^−^ medium ± salts as indicated in figure legends. Glucose was used in place of maltose in complementation assays and in B31 growth assays. Spot plates were incubated at 30° for 2 days (∼48 h) before photographing. Three biological replicates (three independent colonies from a transformation plate) of each construct were tested and spot plate experiments were replicated at least twice. CY162 yeast (Ko and Gaber, 1991) (incapable of K^+^ uptake) growth assays were performed as above using AP media (Rodríguez-Navarro and Ramos, 1984) with 2% glucose and indicated concentrations of KCl and grown at 30°C for 6 days. Media lacking uracil and tryptophan was used in yeast co-expression experiments and the pDR196T vector used for tryptophan selection (Milne et al., 2019). Yeast [^14^C]glucose uptake experiments were performed as described (Milne et al., 2017).

### Field rust resistance screening of the *Lr67* transmembrane region 4 variant

Naturally occurring *Lr67* variants containing only the transmembrane region 4 SNP, encoding the G144R mutation, were crossed from the progenitor donor wheat landrace, AUS4793 to the susceptible cultivar, Avocet S. Homozygous F3 sib selections with G144R were grown in the field alongside Avocet S (Lr67sus) and Avocet+Lr67res (G144R, V387L), inoculated with stripe rust and disease severity assessed on flag leaves of adult plants in accordance with previous phenotyping protocols (Moore et al., 2015).

### Cabinet-grown wheat NaCl treatment

Wheat near-isogenic lines of cv. Thatcher (which carries the wild-type *Lr67sus* allele) and Thatcher+Lr67res (Dyck and Samborski, 1979; Dyck et al., 1994) were subjected to salt treatment at the adult stage of development. Two plants were grown per 15 cm diameter pot in a soil mixture of 50% compost, 25% sand and 25% perlite under an 18-h day/6-h night light cycle at temperatures of 22°C/18°C respectively. Six pots of each genotype were randomly assigned for control or salt-treatment (12 plants per treatment per genotype). All plants were watered with one-quarter strength Hoaglands solution (Hoagland Modified Basal Salt Mixture, Phytotech Labs, Lenexa, KS, USA) one week prior to salt treatment. Half the pots were then treated with 25 mM NaCl + 1.67 mM CaCl_2_, Ca^2+^ requirement as described (Cramer, 2002) in half strength Hoaglands solution at anthesis, and then with 50 mM NaCl + 2.5 mM CaCl_2_ half strength Hoaglands solution six days later whilst control plants were treated with half strength Hoaglands solution only. For each treatment, 350 mL of nutrient solution was applied per pot, and pots were flushed with approximately 700 mL of water 24 hours after treatment. Plants were watered to field capacity 24 h prior to treatment. Flag leaves and penultimate leaves were photographed 11 days after initial NaCl treatment.

### Production and rust inoculation of transgenic barley

A modified version of the *HvSTP13* genomic sequence (HORVU4Hr1G067450.1) incorporating G144R and V387L mutations was synthesized by Epoch Biolabs (Sugar Land, TX, USA), including 1503 bp promoter sequence upstream of the start codon, 1660 bp 3’ UTR sequence after the stop codon and flanked by *NotI* restriction sites. This genomic fragment was inserted into the pVec8 vector and transformed to *Agrobacterium tumifaciens* strain AGL0, which was used to transform barley cv. Golden Promise embryos as described (Harwood, 2014). Transformants were genotyped with a KASPar marker (Table S1) and the KASP Master mix (KBiosciences/LGC Genomics, Teddington, Middlesex, UK).

Plant growth and inoculation was performed as described (Milne et al., 2019). Barley plants were housed in a growth cabinet under 16-h day/8-h night light cycle at a constant temperature of 13°C. After a two-minute heat shock treatment at 42°C, spores of the *P. hordei* pathotype 5457P+ (Singh et al., 2018) (kindly provided by Plant Breeding Institute, Cobbitty, NSW) were mixed with talc powder and sprayed over barley plants. Humidity (∼100%) was maintained by incubating plants in a sealed container at 20°C for 72 h post-inoculation (hpi). Leaves were harvested 8 days post-inoculation, photographed and prepared for microscopic analysis. Microscopic histological assessments were used to determine representative infection site sizes at 8 dpi as described (Ayliffe et al., 2013) by staining chitin present in fungal structures with wheat germ agglutinin conjugated to fluorescein isothiocyanate (WGA-FITC). Microscopic images were photographed using an Axio Imager Z2 microscope and ZEISS ZEN software (Zeiss, North Ryde, NSW, Aust).

## STATISTICAL ANALYSIS

Statistical analysis was performed using Prism software (GraphPad, San Diego, CA, USA) and is described in Figure captions as appropriate. Detailed statistical results are presented in Supplemental Table S3.

## ACCESSION NUMBERS

NCBI accession numbers of sequences used in this study are listed unless specified. *Lr67sus*, MV144992.1; *Lr67res*, KR604817.2; *AtHKT1*, NM_117099.6; *AtSTP13*, AJ344338.1; *GLUT1*, NM_006516.4; *HvSTP13*, MK409638.1; *HvSTP13* genomic clone HORVU4Hr1G067450.1/HORVU.MOREX.r3.4HG0396130 (Ensembl Plants); *Lr34sus*, HL100988.1; *Lr34res*, XM_044586927.1; *SbSTP13*, XM_002465591.2; *TmCLC-0*, X56758.1; *XylE*, AAA79016.1 (European Nucleotide Archive).

## SUPPLEMENTAL DATA

**Supplemental Figure S1**. Effects of external pH change on *X. laevis* oocytes.

**Supplemental Figure S2**. Characterizing the Lr67res gain-of-function in yeast.

**Supplemental Figure S3**. The effect of HvSTP13 site-directed mutants on NaCl sensitivity in yeast.

**Supplemental Figure S4**. NaCl induced leaf tip necrosis.

**Supplemental Table S1**. Primers used in this study.

**Supplemental Table S2**. Amplified and synthesized fragments and constructs used in this study.

**Supplemental Table S3:** Statistical analysis used in this study.

## ACKNOWLEDGEMENTS

The authors wish to thank Dr Sam Henderson for kindly providing B31 and CY162 yeasts; Dr Crystal Wu for kindly providing *TmClC-0* cRNA; Dr Davinder Singh for kindly providing the *P. hordei* isolate 5457P+; Terese Richardson for generating transgenic barley plants; Dr Peter Dodds, Dr Rana Munns, Dr Richard James, Prof. Matthew Gilliham, Dr Caitlin Byrt, Dr Sunita Ramesh, Dr Megan Shelden, Dr Stefanie Wege, Dr Jiaen Qiu, Dr Samantha McGaughey, Wendy Sullivan, and Alexander Taylor for useful discussions and/or technical assistance. We thank Prof. Wolf Frommer for reviewing an earlier draft of this paper. This work was supported by a CSIRO Research Plus Postdoctoral Fellowship (R.J.M.), Bill and Melinda Gates Foundation Grant OPP1131636 (K.E.D., J.Z., E.S.L.), Discovery Early Career Researcher Award DE170100346 (J.B.) and The Australian Research Council also supported this research through CE140100008 (S.D.T.).

## AUTHOR CONTRIBUTIONS

RJM and KED contributed to all experimental work and drafted manuscript. RJM, KED, AR conducted yeast experiments. RJM, KED, JB conducted oocyte experiments. WS, ESL conducted crossing and screening of Lr67 wheat variants. RJM, KED conducted wheat salt treatment experiment. JZ performed transgenic barley rust inoculation and analysis. RJM, KED, JB, ARA, PRR, SDT, ESL conceived experimental plans. All authors commented on draft and approved manuscript.

